# Extracellular adenosine signaling reverses the age-driven decline in the ability of neutrophils to kill *S. pneumoniae*

**DOI:** 10.1101/2020.04.14.041418

**Authors:** Manmeet Bhalla, Shaunna R. Simmons, Alexsandra Abamonte, Sydney E. Herring, Sara E. Roggensack, Elsa N. Bou Ghanem

## Abstract

The elderly are susceptible to serious infections by *Streptococcus pneumoniae* (pneumococcus), which calls for a better understanding of the pathways driving the decline in host defense in aging. We previously found that extracellular adenosine (EAD) shaped polymorphonuclear cell (PMN) responses, which are crucial for controlling infection. EAD is produced by CD39 and CD73, and signals via A1, A2A, A2B and A3 receptors. The objective of this study was to explore the age-driven changes in the EAD pathway and its impact on PMN function. We found in comparison to young mice, PMNs from old mice expressed significantly less CD73, but similar levels of CD39 and adenosine receptors. PMNs from old mice failed to efficiently kill pneumococci *ex vivo*; however, supplementation with adenosine rescued this defect. Importantly, transfer of PMNs expressing CD73 from young mice reversed the susceptibility of old mice to pneumococcal infection. To identify which adenosine receptor(s) is involved, we used specific agonists and inhibitors. We found that A1 receptor signaling was crucial for PMN function as inhibition or genetic ablation of A1 impaired the ability of PMNs from young mice to kill pneumococci. Importantly, activation of A1 receptors rescued the age-associated defect in PMN function. In exploring mechanisms, we found that PMNs from old mice failed to efficiently kill engulfed pneumococci and that A1 receptor controlled intracellular killing. In summary, targeting the EAD pathway reverses the age-driven decline in PMN antimicrobial function, which has serious implications in combating infections.

## Introduction

Despite the availability of vaccines and antibiotics, *S. pneumoniae* remain the leading cause of community-acquired pneumonia in the elderly (Henig & Kaye, 2017). In recent Active Bacterial Core surveillance reports, people above 50 accounted for 71% of all pneumococcal cases and 82% of associated deaths (CDC, 2017). Immunosenescence, the overall decline in immunity that occurs with age, contributes to the increased susceptibility of the elderly to infection (Krone, van de Groep, Trzcinski, Sanders, & Bogaert, 2014). We and others previously found that neutrophils (polymorphonuclear leukocytes, or PMNs) are required for host defense against *S. pneumoniae* infections (Bou Ghanem et al., 2015; Hahn et al., 2011) as they are needed for initial control of bacterial numbers upon infection (Bou Ghanem et al., 2015). PMN anti-microbial function is known to be dysregulated with aging. There are reports of decreased phagocytic capacity, ROS production, extracellular trap formation and overall killing of various pathogens, including *S. pneumoniae* by PMNs from aging hosts (Boe, Boule, & Kovacs, 2017; Hazeldine et al., 2014; Wenisch, Patruta, Daxbock, Krause, & Horl, 2000). However, the host pathways behind this age-driven decline in PMN function remain incompletely elucidated.

Extracellular adenosine (EAD) is key for host resistance to pneumococcal infection (Bou Ghanem et al., 2015). Upon tissue injury triggered by a variety of insults, including infection, ATP is released from cells and metabolized to adenosine by the sequential action of two extracellular enzymes, CD39 that converts ATP to AMP, and CD73 that de-phosphorylates AMP to EAD (Thompson et al., 2004). Conversely, EAD is broken down by adenosine deaminase (ADA). We previously found that EAD production by CD73 was crucial for host resistance against *S. pneumoniae* lung infection in mice. Mice that had CD73 pharmacologically inhibited or genetically ablated suffered dramatically higher pulmonary bacterial numbers, systemic spread of the infection and increased lethality upon *S. pneumoniae* lung infection (Bou Ghanem et al., 2015). Importantly, CD73 controlled PMN anti-microbial phenotype against *S. pneumoniae* (Siwapornchai et al., 2020). CD73 expression and EAD production by PMNs was required for the ability of these cells to kill and clear *S. pneumoniae* (Siwapornchai et al., 2020; Thompson et al., 2004). EAD is recognized by four G-protein coupled receptors, A1, A2A, A2B and A3 (Hasko, Linden, Cronstein, & Pacher, 2008). These receptors ubiquitously expressed on many cell types including PMNs (Barletta, Ley, & Mehrad, 2012) and can have opposing effects on immune responses (Kumar & Sharma, 2009). The adenosine receptor(s) mediating the anti-microbial activity of PMNs against *S. pneumoniae* remain unknown.

Aging is accompanied by changes in EAD production and signaling (Crosti et al., 1987; Headrick, 1996; Mackiewicz et al., 2006; Willems, Ashton, & Headrick, 2005). For instance, changes in the EAD pathway contribute to the age-related decline of brain (Mackiewicz et al., 2006), metabolic (Rolband, Furth, Staddon, Rogus, & Goldberg, 1990) and cardiac function (Willems et al., 2005). However, the role of the extracellular adenosine pathway in immunosenescence remains practically unexplored. One study reported that aging resulted in changes in T cell production and responsiveness to EAD and inhibiting A2A receptor reversed age-related deficiencies in chemotaxis, proliferation and cytokine production by T cells (Hesdorffer et al., 2012). We previously found that triggering A1 receptor signaling in old mice significantly enhanced their resistance to pneumococcal lung infection and reduced the ability of *S. pneumoniae* to bind pulmonary epithelial cells (Bhalla et al., 2020). In this study, we explored the age-driven changes in the expression of each of the EAD pathway enzymes and receptors on PMN. We identified for the first time a key role for adenosine production and signaling in the age-associated decline in anti-microbial function of PMNs.

## Results

### The ability of PMNs to kill *S. pneumoniae* is impaired with aging

Aging is accompanied by increased susceptibility to *S. pneumoniae* infection and we previously found that PMNs are required for host protection against these bacteria. To test if PMN function declines with age, we compared *ex vivo* opsonophagocytic killing of pneumococci by bone-marrow derived PMNs isolated from young (2 months) vs. old mice (18-22 months). We found that the anti-microbial activity of PMNs was significantly impaired with age, where we observed a 5-fold decline in the ability of PMNs from old mice to kill *S. pneumoniae* as compared to young controls (Fig 1). This is in line with previous studies highlighting the decline in PMN function with age (Simell et al., 2011).

**Figure 1.**
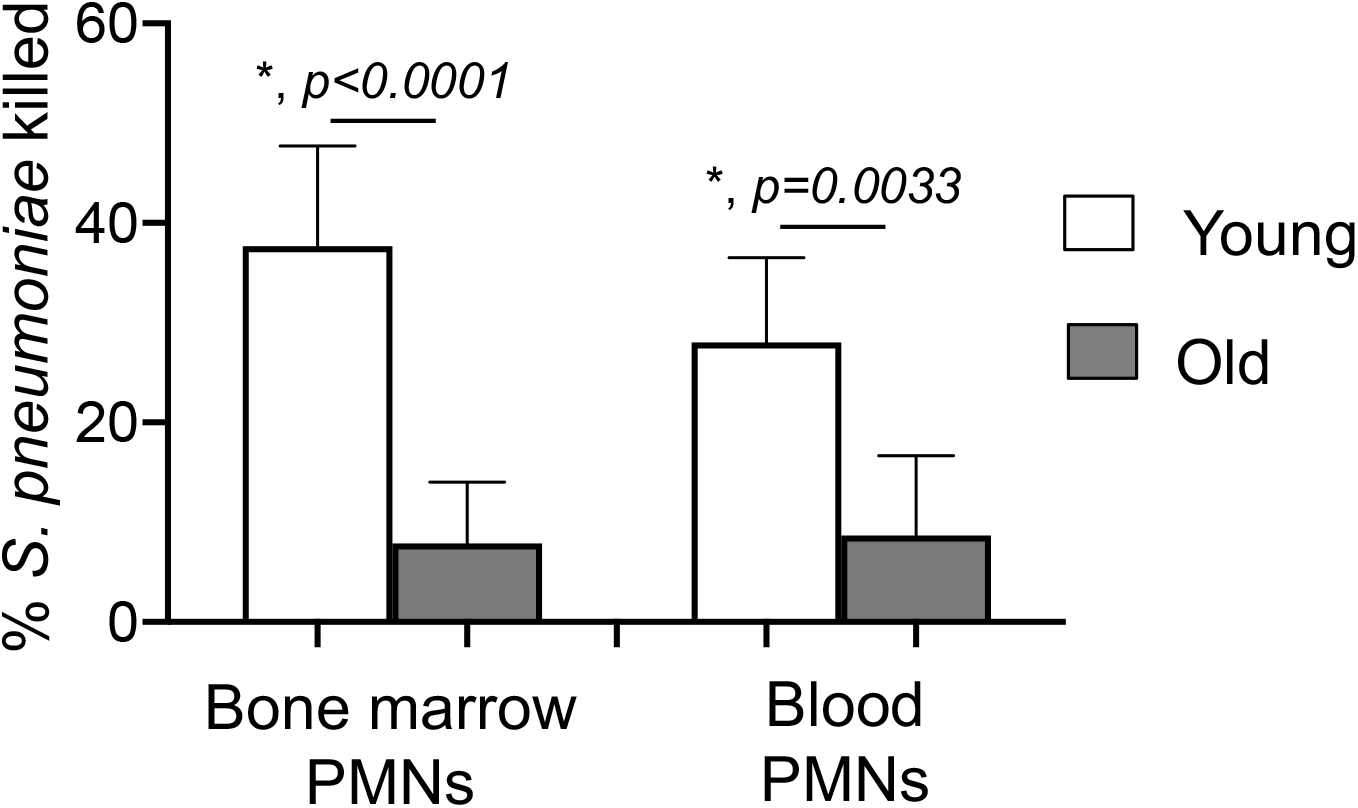
PMNs from old mice fail to efficiently kill *S. pneumoniae.* (A) PMNs were isolated either from the bone marrow or blood of young (2 month) and old (18-22 month) C57BL/6 mice and infected with *S. pneumoniae* pre-opsonized with homologous sera for 45 minutes at 37°C. Reactions were stopped on ice and viable CFU were determined after serial dilution and plating. The percentage of bacteria killed upon incubation with PMNs was determined by comparing surviving CFU to a no PMN control. Data shown are pooled from three separate experiments (n=3 biological replicates or mice per strain) where each condition was tested in triplicate (n=3 technical replicates) per experiment. Asterisks indicate significant differences calculated by one-way ANOVA followed by Tukey’s test.

### Aging is accompanied by changes in the expression of EAD-pathway enzymes on PMNs

We previously found that extracellular adenosine (EAD) production by PMNs is required for their anti-microbial activity against *S. pneumoniae* (Siwapornchai et al., 2020). To test if aging is associated with changes in the expression of EAD-producing and degrading enzymes, we compared the expression of CD73, CD39 and ADA on the surface of bone-marrow derived PMNs using flowcytometry. We found that compared to young controls, PMNs from old mice expressed significantly less CD73, the EAD-producing enzyme, but significantly higher levels of the EAD-degrading enzyme ADA, while the expression of CD39 was unchanged (Fig 2A). These findings demonstrate that expression of EAD enzymes on PMNs are significantly altered with age.

**Figure 2.**
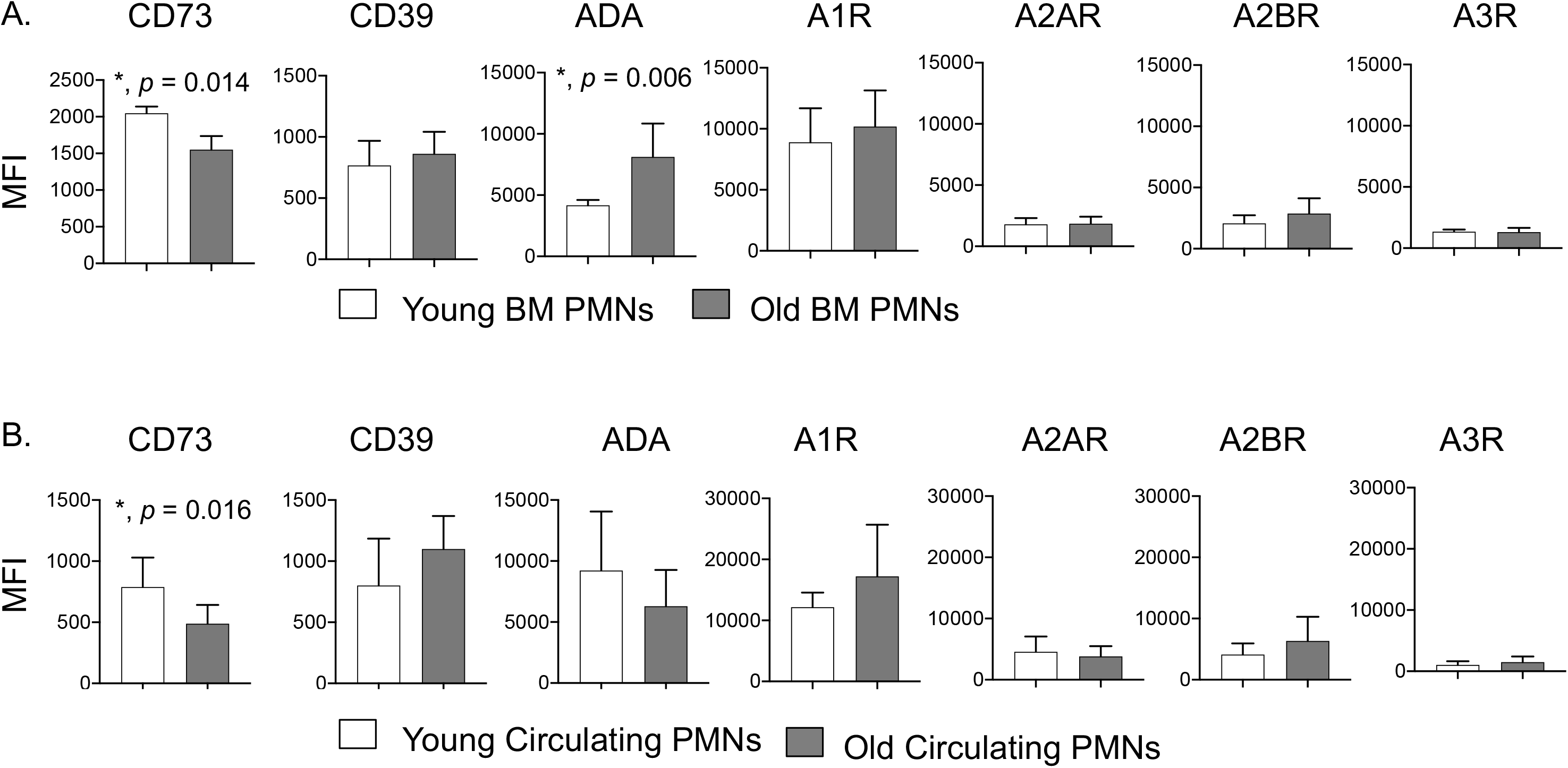
Expression of EAD-pathway enzymes on PMNs are altered with age. (A) Bone marrow cells were isolated from young and old C57BL/6 mice and expression of EAD-pathway components was assessed by flow cytometry. (B) Blood was collected from young and old C57BL/6 mice and expression of EAD-pathway components was assessed by flow cytometry. (A-B) We gated on PMNs (Ly6G^+^CD11b^+^ cells) and measured the expression (mean fluorescent intensity or MFI) of CD73, CD39, ADA, A1R, A2AR, A2BR and A3R. (A) Data shown are pooled from three separate experiments (n=3 mice per age group) with each condition tested in triplicate. (B) Data are pooled from four separate experiments with 7 mice per age group. Asterisks indicate significant differences from uninfected controls calculated by Student’s t-test.

### CD73 expression on circulating PMNs decreases with age

Next, we wanted to confirm the importance of our findings *in vivo.* We compared the expression of the EAD-pathway components on circulating PMNs in young and old mice. We found that similar to what we observed in bone-marrow derived cells, circulating PMNs in old mice expressed significantly lower levels of CD73 on their surface (Fig 2B). We did not find any differences in the expression of the rest of the EAD pathway enzymes (CD39, ADA) on circulating PMNs (Fig 2B). Further, circulating PMNs in old mice also failed to efficiently kill *S. pneumoniae* (Fig 1). These findings confirm that there are age-driven changes in CD73 expression on PMNs circulating *in vivo*.

### Adoptive transfer of PMNs from wild type but not CD73^−/−^ young mice boosts resistance of old hosts to *S. pneumoniae*

To test the importance of CD73 expression by PMNs in host resistance against *S. pneumoniae,* we adoptively transferred 2.5×10^6^ PMNs isolated from the bone marrow of WT or CD73^−/−^ young mice into old mice. We then infected mice one hour later with 5×10^5^ CFU of *S. pneumoniae* i.t. and compared clinical scores and bacterial burdens in different organs at 18 hours post infection. We found that transfer of WT PMNs from young mice significantly reduced pulmonary bacterial burdens in old mice (Fig 3A). WT PMNs further reduced the systemic spread of pneumococci resulting in a 50-fold reduction in blood and brain bacterial loads when compared to no transfer controls (Fig 3B and C). Importantly, transfer of WT PMNs from young mice ameliorated clinical signs of the disease in aged hosts (Fig 3D). In contrast, transfer of PMNs from CD73^−/−^ young mice had no effect on reducing bacterial burdens or improving the disease score in old recipients (Fig 3). These findings demonstrate that PMNs from young mice are sufficient to boost resistance of old hosts to *S. pneumoniae* infection and that this protection is dependent on CD73 expression by PMNs.

**Figure 3.**
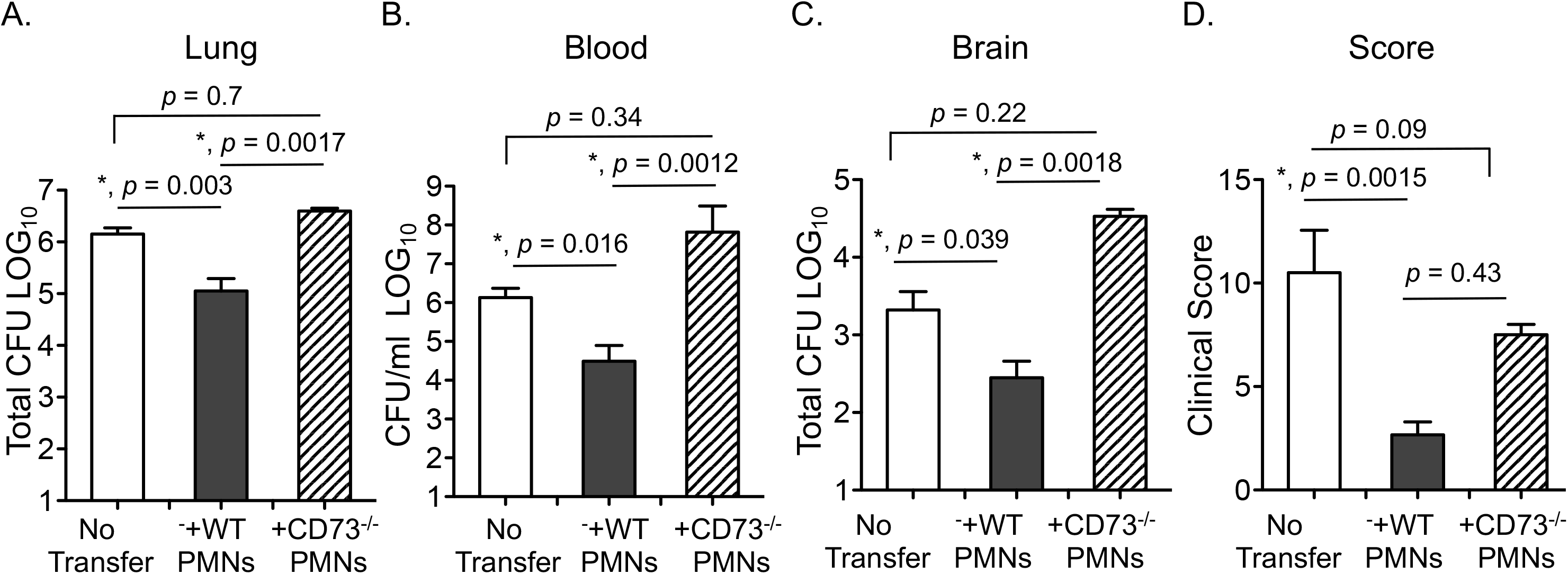
Adoptive transfer of PMNs from young wild type but not CD73^−/−^ mice boosts resistance of old mice to *S. pneumoniae.* Old male C57BL/6 mice were mock treated (no transfer) or adoptively transferred 2.5×10^6^ of the indicated PMNs isolated from the bone marrow of young male C57BL/6 or CD73^−/−^ mice. One hour post transfer, mice were infected i.t with 5×10^5^ CFU of *S. pneumoniae* and bacterial numbers in the lung (A), blood (B) and brain (C) as well as (D) clinical scores were determined 18 hours post infection. Asterisks indicate significant differences calculated by one-way ANOVA followed by Tukey’s test. Pooled data from three separate experiments with n=8 mice no transfer group, n=8 mice +WT PMNs and n=5 mice + CD73^−/−^ PMNs are shown.

### Supplementation with adenosine reverses the age-driven decline in PMN antimicrobial function

Our above findings suggested that the ability of PMNs to produce EAD declines with age. As circulating and bone marrow-derived PMNs exhibited mostly similar phenotypes (Fig 1 and 2), for feasibility, we proceeded with experiments using PMNs isolated from the bone marrow that are routinely used for *ex vivo* assays (Bou Ghanem et al., 2015; Siwapornchai et al., 2020; Standish & Weiser, 2009). To directly determine if EAD production differs with age, we measured the baseline levels of EAD produced by PMNs within 5 minutes of culture in the *ex vivo* assay prior to infection. To differentiate how much of EAD was CD73-dependent, we measured the amounts produced by CD73^−/−^ PMNs. We found that the amount of EAD produced by PMNs from old wildtype mice was comparable to those produced by CD73^−/−^ PMNs (Fig 4A). In contrast, PMNs from young wildtype mice produced 3-fold more EAD as compared to PMNs from either old or CD73^−/−^ mice (Fig 4A). Upon pneumococcal infection, we observed a 6-fold increase in EAD produced by PMNs from young mice, but no significant increase in PMN cultures from old mice (Fig 4B). To rule out EAD release from dying cells as a potential source for the observed differences between PMNs from old and young mice, we compared apoptosis and necrosis using Annexin V/PI staining as previously described (Siwapornchai et al., 2020). We found no difference in the number of apoptotic/necrotic PMNs between the different mice groups (not shown), indicating that the differences in EAD production is not due to differences in cell viability.

**Figure 4.**
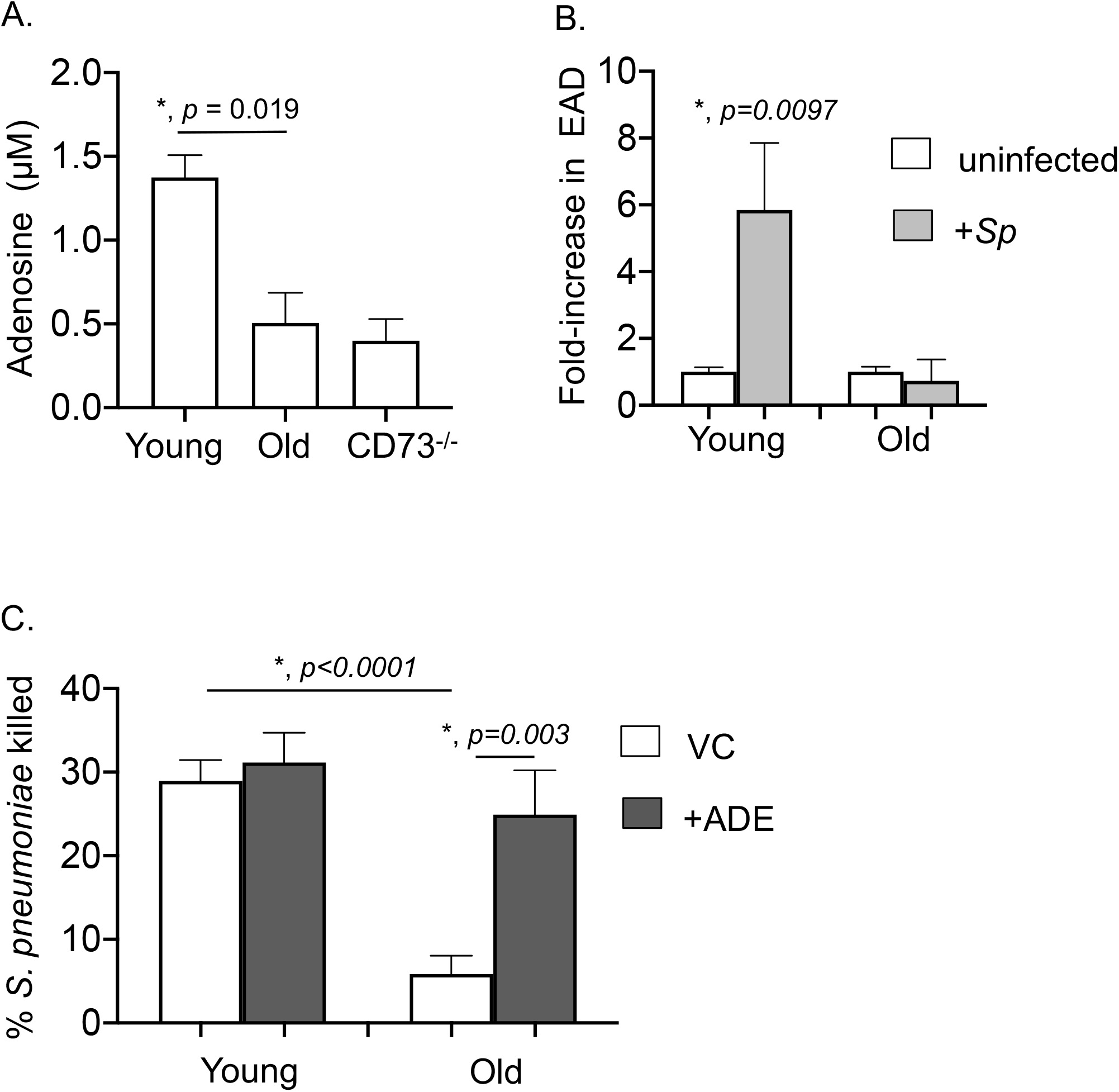
Extracellular adenosine rescues the ability of PMNs from old mice to kill *S. pneumoniae.* (A) PMNs were isolated from the bone marrow of young and old C57BL/6 mice as well as CD73^−/−^ young mice and incubated in assay buffer for 5 minutes. The amount of adenosine in the supernatants was then measured. Asterisks indicate significant differences calculated by Student’s test. (B) Bone marrow PMNs isolated from young and old C57BL/6 mice were infected with *S. pneumoniae* pre-opsonized with homologous sera for 45 minutes at 37°C. The amount of adenosine in the supernatants was then measured and the fold-change in extracellular adenosine production was calculated by dividing the values of infected reactions by uninfected controls for each condition. Asterisks indicate significantly different from 1 by one-sample t-test. (A-B) are representative data from one of three separate experiments, where each condition was tested in quadruplicate. (C) PMNs were isolated from the bone marrow of young and old C57BL/6 mice and treated with 1μM Adenosine (+ADE) or PBS (VC) for 30 minutes at 37°C. The reactions were then infected with *S. pneumoniae* pre-opsonized with homologous sera for 45 minutes at 37°C. Reactions were stopped on ice and viable CFU were determined after serial dilution and plating. The percentage of bacteria killed upon incubation with PMNs was determined by comparing surviving CFU to a no PMN control. Data shown are pooled from three separate experiments (n=3 biological replicates or mice per strain) where each condition was tested in triplicate (n=3 technical replicates) per experiment. Asterisks indicate significant differences calculated by one-way ANOVA followed by Tukey’s test.

To test if supplementing with EAD rescues the antimicrobial function of PMNs from old mice, we added EAD to the opsonophagocytic reactions. Strikingly, we found that supplementation with 1μM EAD fully restored the ability of PMNs from old mice to kill bacteria to a comparable level observed with young controls (Fig 4C). Taken together, these findings suggest that the age-driven impairment in pneumococcal killing is largely due to a decrease in EAD production, and that this impairment can be fully reversed *in vitro* by supplementation with EAD.

### A1 receptor signaling is required for the ability of PMNs to kill *S. pneumoniae*

EAD can signal via four G-protein coupled receptors A1, A2A, A2B and A3, which are ubiquitously expressed (Hasko et al., 2008). Therefore, we wanted to pinpoint which adenosine receptor(s) was required for PMN anti-pneumococcal function. First, we compared the expression of the different adenosine receptors by flowcytometry as previously described (Bhalla et al., 2020). We found that the A1 receptor was more highly expressed on PMNs as compared to the other receptors, while A3 expression was the lowest (Supplemental Fig 1A). As the A1 receptor antibody is polyclonal, we further confirmed A1 receptor expression on PMNs by western blots comparing wildtype (WT) and A1 receptor knock-out (A1R^−/−^) mice (Supplemental Fig 1B). We did not observe any differences in the expression of any of the adenosine receptors on bone-marrow derived or circulating PMNs with age (Fig 2A and B).

To test which of the adenosine receptors were important for PMN antimicrobial function, we treated PMNs from young mice with specific inhibitors for each of the different receptors and compared their ability to kill bacteria *ex vivo*. We found that blocking A1 receptor signaling significantly blunted the ability of PMNs from young mice to kill *S. pneumoniae* (Fig 5A), while inhibition of the other three receptors did not significantly impair bacterial killing by PMNs. Importantly, none of the adenosine receptor inhibitors had a direct effect on bacterial viability (Supplemental Fig 2), indicating that effect on bacterial killing was specifically due to PMN anti-bacterial function. We further confirmed the role of A1 receptor in the anti-bacterial function of PMNs by comparing the ability of WT, A1R^−/+^ and A1R^−/−^ PMNs to kill bacteria *ex vivo.* We found that the ability of A1R^−/+^ PMNs to kill pneumococci was severely impaired as compared to WT controls, while A1R^−/−^ PMNs completely failed to kill *S. pneumoniae* (Fig 5B). In fact, bacterial numbers increased in the presence of A1R^−/−^ PMNs (Fig 5B). Taken together, these findings demonstrate that A1 receptor signaling is required for the anti-pneumococcal activity of PMNs.

**Figure 5.**
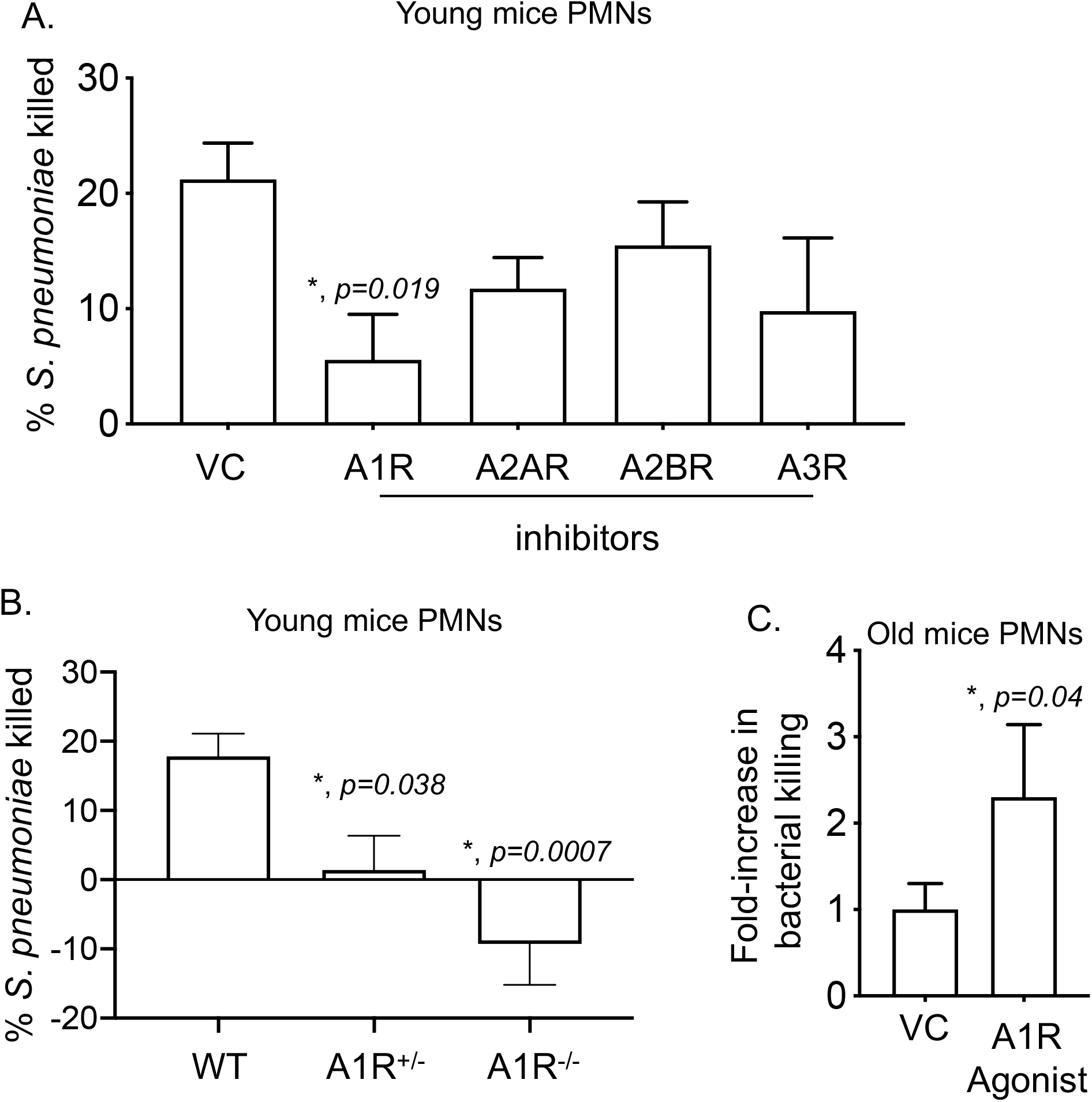
A1 receptor signaling is required for the ability of PMNs to kill *S. pneumoniae.* (A) PMNs isolated from the bone marrow of young C57BL/6 mice were treated with either the A1 receptor inhibitor 8-Cyclopentyl-1-3-dipropylxanthine (3.9nM), A2A receptor inhibitor 3,7-Dimethyl 1-1-propargylxanthine (11 μM), A2B receptor inhibitor MRS 1754 (1.97nM), A3 receptor inhibitor MRS 1191 (92 nM) or PBS as vehicle control (VC) for 30 minutes at 37°C. (B) PMNs isolated from the bone marrow of young wildtype (WT), A1R^−/+^ and A1R^−/−^ mice. (C) PMNs isolated from the bone marrow of old C57BL/6 mice were treated with either the A1 receptor agonist 2-Chloro-N6-Cyclopentyl adenosine (0.8 nM), Adenosine (1μM) or PBS as vehicle control (VC) for 30 minutes at 37°C. (A-C) PMNs were then infected with pre-opsonized *S. pneumoniae* for 45 minutes at 37°C. Reactions were plated on blood agar plates and the percentage of bacteria killed compared to a no PMN control under the same conditions was calculated. (C) The fold-change in bacterial killing was then calculated by dividing the values of A1 receptor agonist treated reactions by vehicle treated controls for each condition. (A-B) Data shown are pooled from three separate experiments (n=3 mice per strain) with each condition tested in triplicate. Asterisks indicate significant differences vs. VC reactions calculated by one-way ANOVA followed by Dunnet’s test. (C) Data shown are pooled from four separate experiments (n=4 mice per strain). Asterisks indicate significantly different from 1 by one-sample t-test.

### Activation of A1 receptor signaling rescues the ability of PMNs from old mice to kill *S. pneumoniae*

Next we wanted to test if A1 receptor signaling plays a role in PMN function in old mice. To test this, we treated PMNs from old mice with 2-Chloro-N6-Cyclopentyl adenosine, an A1 receptor specific agonist, whose specificity we had confirmed before (Bhalla et al., 2020). We then measured the ability of PMNs to kill bacteria *ex vivo*. We found that in comparison to vehicle control, treatment with the A1 receptor agonist boosted the ability of PMNs from old mice to kill *S. pneumoniae* more than two-fold (Fig 5C). This demonstrates that activation of A1 receptor signaling rescues the age-associated defect in PMN function.

### ROS production and release of MPO and CRAMP are not impaired with aging

We then wanted to explore why PMNs from old mice fail to efficiently kill *S. pneumoniae.* We previously found that CD73 was important for optimal ROS production by PMNs which contributes to their ability to kill bacteria (Siwapornchai et al., 2020). However, we found no differences in the ability of PMNs from young and old mice to produce intracellular or extracellular ROS in response to *S. pneumoniae* (Supplemental Fig 3A and B). Previous work has also showed that antimicrobial peptides and enzymes can directly kill pneumococci (Habets, Rozen, & Brockhurst, 2012; Xiang et al., 2017). Therefore, we measured levels of Cathelin-Related Anti-Microbial Peptide (CRAMP) and Myeloperoxidase (MPO) released by PMNs. Again, we found no significant differences in the amount of MPO or CRAMP (Supplemental Fig 3C and D) in the supernatants of PMNs from young vs. old hosts. Thus, the age-driven decline in anti-pneumococcal PMN responses do not appear to derive from ROS production or release of MPO and CRAMP.

### Aging is accompanied by a defect in intracellular killing of engulfed bacteria

Bacterial uptake was previously demonstrated to be important for killing by PMNs (Standish & Weiser, 2009), therefore, we wanted to test if phagocytosis of pneumococci was impaired with aging. To do so, we established a flowcytometry-based bacterial uptake assay with GFP-expressing bacteria and inside-out staining (Smirnov, Solga, Lannigan, & Criss, 2015). We first infected PMNs with GFP-expressing *S. pneumoniae* and differentiated between associated vs. engulfed bacteria by staining the cells with anti *S. pneumoniae* antibodies (PE-labeled). We found that of all bacteria that associated with PMNs within 15 minutes, 40% of them were engulfed (GFP^+^/ PE^−^) (Supplemental Fig 4A and B). We confirmed the validity of the assay using Cytochalasin D which impairs phagocytosis (Standish & Weiser, 2009) and found that in its presence, the majority of PMN-associated bacteria (~90%) remained extracellular (Supplemental Fig 4A and B). When we compared hosts across age, we found no significant differences in bacterial association or uptake between PMNs from young or old mice (Supplemental Fig 4B and C). However, when we compared the amounts of engulfed bacteria that had survived using a gentamicin-protection assays we had previously established (Siwapornchai et al., 2020), we found that there were four-fold more viable bacteria in PMNs from old mice as compared to young counterparts (Fig 6A). These findings demonstrate that while phagocytosis of *S. pneumoniae* does not change with age, the ability of PMNs to kill the engulfed bacteria is significantly diminished in PMNs from old mice compared to young mice.

**Figure 6.**
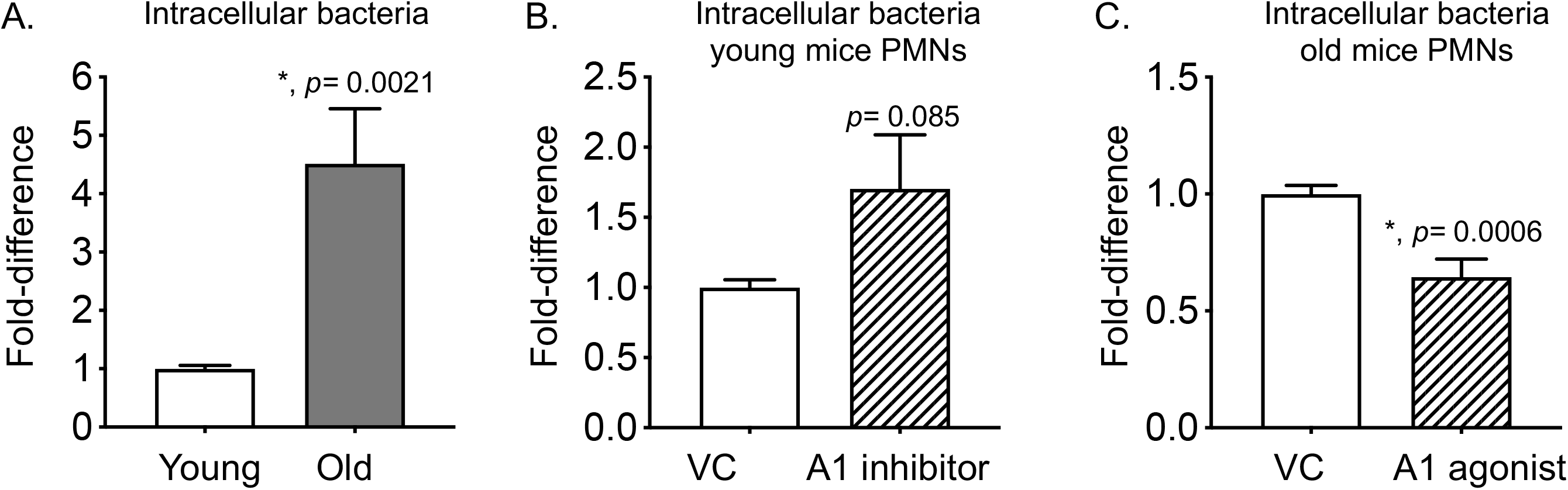
The ability of PMNs to kill intracellular bacteria declines with age and is controlled by A1 receptor signaling. (A) PMNs were isolated from the bone marrow of young and old male C57BL/6 mice. (B) PMNs isolated from the bone marrow of young mice were treated with either the A1 receptor inhibitor 8-Cyclopentyl-1-3-dipropylxanthine (3.9nM) or PBS as vehicle control (VC) for 30 minutes at 37°C. (C) PMNs isolated from the bone marrow of old mice were treated with either the A1 receptor agonist 2-Chloro-N6-Cyclopentyl adenosine (0.8 nM) or PBS as vehicle control (VC) for 30 minutes at 37°C. (A-C) PMNs were then infected with pre-opsonized *S. pneumoniae* for 15 minutes at 37°C. Gentamicin (100μg/ml) was then added for 30 minutes to kill extracellular bacteria. PMNs were washed and plated on blood agar plates to determine the % of viable bacteria of the original infecting inoculum. The fold-difference in viable intracellular bacteria was then calculated by dividing the values of (A) old reactions by young controls, (B) A1 receptor inhibitor treated reactions by vehicle treated controls and (C) A1 receptor agonist treated reactions by vehicle treated controls. Asterisks indicate significantly different from 1 by one-sample t-test.

### A1 receptor signaling controls intracellular killing of engulfed *S. pneumoniae*

Since intracellular killing of bacteria was impaired in aging, we wanted to determine whether this was controlled by A1 receptor signaling. To test that, we treated PMNs from young mice with the A1 receptor inhibitor and compared intracellular survival of *S. pneumoniae.* We found that upon inhibition of A1 receptor signaling, the number of viable intracellular bacteria in young PMNs almost doubled (Fig 6B). On the other hand, treatment of PMNs from old mice with the A1 receptor agonist significantly reduced the amounts of intracellular bacteria that survived by half (Fig 6C). The observed differences in bacterial viability were not due to any effect on bacterial uptake by PMNs, as inhibition or activation of A1 receptor had no effect on bacterial phagocytosis by PMNs from young and old mice, respectively (Supplemental Fig 4B and C). Taken together, these data demonstrate that A1 receptor signaling significantly contributes to the ability of young PMNs to kill engulfed pneumococci and that this process is impaired with aging. Importantly, activating A1 receptor signaling restores the ability of PMNs from old mice to kill engulfed *S. pneumoniae*.

## Discussion

Extracellular adenosine is known to regulate PMN function (Barletta et al., 2012) and more recently, the role of this pathway in host resistance against pulmonary infections has been also highlighted (Lee & Yilmaz, 2018). However, the role of EAD in immunosenescence remains largely unexplored. A few studies previously tracked the expression of EAD producing enzymes on T cells in aging and found that in mice, expression of CD39 and CD73 were upregulated on splenic regulatory T cells (Alam, Cavanaugh, Pereira, Babu, & Williams, 2020). Similarly, activated CD4^+^T cells from elderly human volunteers expressed higher levels of CD39 than those from younger donors (Fang et al., 2016). In contrast, the expression of CD73 was lower on CD8^+^T cells in elderly healthy humans as compared to younger controls (Jeske et al., 2020). Whether the expression of EAD-pathway components on PMNs is altered with aging remained completely unexplored until now. Here, we found that expression of CD73 on PMNs significantly declined with aging, while levels of CD39 were unchanged. We also found that surface expression of ADA, the enzyme that breaks down adenosine, was actually higher in bone-marrow derived PMNs in old mice as compared to young controls. The activity and levels of ADA were previously shown to decline in peripheral blood lymphocytes of elderly donors (Crosti et al., 1987) and in senescent CD8^+^T cells (Parish et al., 2010). To our knowledge, this is the first study to demonstrate that aging is accompanied by changes in EAD-pathway enzymes on PMNs.

The changes in EAD-pathway enzymes in aged mice were associated with lower amounts of adenosine present in the extracellular milieu of PMN cultures from old mice as compared to young counterparts. EAD can be either released into the extracellular environment from intracellular compartments via equilibrative nucleoside transporters (Boswell-Casteel & Hays, 2017), or can be produced as a breakdown product of extracellular ATP, a process that requires CD39 and CD73 (Eltzschig, Macmanus, & Colgan, 2008). At baseline the amount of EAD produced by PMNs from old mice was comparable to that made by CD73^−/−^ PMNs, suggesting the amounts released were CD73-independent. However, PMNs from young mice, which expressed higher levels of CD73, produced significantly more EAD even at baseline. Upon infection, we further observed a significant increase in EAD production, but only by PMNs from young mice. Pneumolysin, a toxin produced by *S. pneumoniae* (Marriott, Mitchell, & Dockrell, 2008), triggers the release of extracellular ATP from infected cells (Domon et al., 2016; Hoegen et al., 2011). Therefore, it is likely that in PMNs from young mice, the increase in EAD production seen upon infection is due to the conversion of ATP into EAD by the sequential action of CD39 and CD73, a process that is impaired in PMNs from old mice due to blunted expression of CD73.

We previously found that CD73 expression and autocrine EAD production by PMNs is crucial for the antimicrobial phenotype of PMNs during *S. pneumoniae* infection (Siwapornchai et al., 2020). EAD can signal via A1, A2A, A2B and A3 receptors (Hasko et al., 2008). As these receptors can have opposing effects on immune responses (Kumar & Sharma, 2009), with some of them, in particular the higher affinity ones such as A2A and A2B being suppressive in most cases (Barletta et al., 2012), it was important to identify which receptor was mediating the antimicrobial activity of PMNs. Adenosine binds its receptors with different affinities ranging from an EC_50_ <0.5μM for the high affinity receptors A1 and A3, to an EC_50_ >0.6μM for the intermediate affinity receptor A2A and an EC_50_ between 16-64μM for the low affinity A2B receptor (Hasko et al., 2008). Here, we found that supplementation of PMNs with 1μM of adenosine was sufficient to restore the antimicrobial function of PMNs from old mice suggesting that the higher affinity receptors were involved. In fact, using a combination of pharmacological and genetic approaches, we pinpointed A1 receptor as the receptor required for the ability of PMNs to kill *S. pneumoniae.* Importantly, triggering this receptor reversed the age driven defects in the ability of PMNs to kill these bacteria. This is in line with our previous findings demonstrating that activating A1 receptor signaling *in vivo* boosts the resistance of old mice to pneumococcal pneumonia by enhancing pulmonary bacterial clearance and reducing systemic spread of the infection (Bhalla et al., 2020).

In exploring mechanisms of why PMN function was impaired with aging we examined ROS production, which we and others have previously reported to be controlled by adenosine (Cronstein, Daguma, Nichols, Hutchison, & Williams, 1990; Salmon & Cronstein, 1990; Siwapornchai et al., 2020). However, as has been previously reported during pneumococcal infection (Krone, Trzcinski, Zborowski, Sanders, & Bogaert, 2013), we found that ROS production was not altered with aging. Similarly, we found that release of antimicrobial MPO and CRAMP as well as phagocytosis were not impaired in response to *S. pneumoniae* infection. Previously A1 receptor was shown to enhance phagocytosis by PMNs (Salmon & Cronstein, 1990), which we did not observe in our study. However, the previous work was performed using erythrocyte coated beads and stimulation with FMLP or PMA (Cronstein et al., 1990; Salmon & Cronstein, 1990), while here we are performing our studies with live bacteria that can actively modulate PMN responses. Rather, we found that PMNs from old mice failed to efficiently kill engulfed bacteria. The fate of intracellular *S. pneumoniae* and the mechanisms of clearance of these engulfed bacteria have not been fully elucidated. We identify here A1 receptor signaling as the host pathway controlling intracellular killing of pneumococci. Importantly, activation of this receptor enhanced the ability of PMNs from old mice to kill intracellular bacteria.

In conclusion, we demonstrate here for the first time, that immunosenescence of PMNs is shaped by the EAD pathway. We find that the age-driven impairment in bacterial killing by these innate immune cells is largely due to a decrease in EAD production and signaling. Importantly, supplementing with EAD and triggering A1 receptor signaling fully reverses the age-driven decline in PMN antimicrobial function. This has serious implications for considering the future use of clinically available adenosine-based drugs (Jacobson, Tosh, Jain, & Gao, 2019) to combat immunosenescence and pneumococcal pneumonia in the susceptible elderly population.

## Experimental Procedures

### Mice

Young (2 months) and Old (18-22 months) male C57BL/6 mice were purchased from Jackson Laboratories (Bar Harbor, ME) and the National Institute on Aging colonies and housed in a specific-pathogen free facility at the University at Buffalo. CD73^−/−^ mice on C57BL/6 background (Thompson et al., 2004) and B6N.129P-*Adora1*^tm1Bbf^/J mice were purchased from Jackson Laboratories (Bar Harbor, ME), bred at our facility and genotyped as previously described (Johansson et al., 2001). All work was performed in accordance with the recommendations in the Guide for the Care and Use of Laboratory Animals published by the National Institutes of Health. Procedures were reviewed and approved by the University at Buffalo Institutional Animal Care and Use Committee.

### Bacteria

*S. pneumoniae* TIGR4 AC316 strain (serotype 4) (Greene, Narciso, Filipe, & Camilli, 2015) was a kind gift from Andrew Camilli. Bacteria were grown at 37°C in 5% CO_2_ in Todd-Hewitt broth supplemented with 0.5% yeast extract and oxyrase untill mid-exponential phase. Aliquots were frozen at −80°C in the growth media with 20% (v/v) glycerol. Prior to use, aliquots were thawed on ice, washed and diluted in PBS. Bacterial concentrations were confirmed by serial dilution and dribble plating on Tryptic Soy Agar plates supplemented with 5% sheep blood agar.

### Flowcytometry

Whole blood was collected from mice using cardiac puncture using EDTA as an anti-coagulant. Bone marrow cells were harvested from femurs and tibias of mice, flushed with RPMI 1640 supplemented with 10% FBS and 2 mM EDTA, and resuspended in PBS. Red blood cells were removed by treatment with a hypotonic lysis buffer (Lonza). The cells were resuspended in FACS buffer (HBSS/ 1% FBS) then treated with Fc block (anti-mouse clone 2.4G2) and stained with specific antibodies purchased from eBioscience and BD. The following anti-mouse antibodies were used: Ly6G (IA8), CD11b (M1/70), CD39 (24DMS1) and CD73 (TY/11.8). Anti-mouse ADA (ab175310) was purchased from abcam. Staining for adenosine receptor antibodies was done as previously described (Bhalla et al., 2020). Cells were permeabilized using the BD Cytofix/Cytoperm kit. The following unconjugated primary rabbit polyclonal anti-adenosine receptor antibodies were purchased from abcam and used: A2a (ab3461), A2b (ab222901), A3 (ab203298) and A1 (ab82477). Rabbit polyclonal IgG (ab37415) purchased from abcam and used as an isotype control. Secondary PE-conjugated anti-Rabbit IgG was used (12473981; Invitrogen). Fluorescence intensities were measured on a BD FACS Fortessa and data were analyzed using FlowJo.

### PMN isolation

PMNs were isolated from the bone marrow through density gradient centrifugation, using Histopaque 1119 and Histopaque 1077 as previously described (Swamydas & Lionakis, 2013). PMNs were isolated from the circulation through density gradient centrifugation, using a Percoll gradient as previously described (Siwapornchai et al., 2020). The isolated PMNs were resuspended in Hanks’ Balanced Salt Solution (HBSS)/0.1% gelatin without Ca^2+^ and Mg^2+^, and used in subsequent assays. Purity was measured by flow cytometry using CD11b and Ly6G and 85-90% of enriched cells were positive Ly6G/CD11b^+^.

### Adoptive Transfer of PMNs

Bone marrow PMNs were isolated from uninfected mice as described above and 2.5×10^6^ cells were adoptively transferred as previously described (Siwapornchai et al., 2020). Control groups received PBS. One hour following transfer, mice were challenged intra-tracheally with 5×10^5^ CFU of *S. pneumoniae.* Twenty-four hours post infection, mice were scored for clinical signs of the disease ranging from healthy (0) to severely sick (21) as previously described (Bhalla et al., 2020). Mice were euthanized and the lungs, brain, and blood were collected and plated on blood agar plates for CFU.

### Opsonophagocytic (OPH) Killing Assay

The ability of PMNs to kill *S. pneumoniae ex vivo* was measured using a well-established Opsonophagocytic (OPH) killing assay as previously described (Bou Ghanem et al., 2015; Lysenko et al., 2007; Siwapornchai et al., 2020). Briefly, 2×10^5^ PMNs were incubated with 1×10^3^ bacteria grown to mid log phase and pre-opsonized with 3% mouse sera in 100μl reactions of HBSS/0.1% gelatin. Reactions were rotated for 45 minutes at 37°C. Where indicated, PMNs were incubated with adenosine (100μM), A1 receptor inhibitor 8-Cyclopentyl-1-3-dipropylxanthine (3.9nM), A2A receptor inhibitor 3,7-Dimethyl 1-1-propargylxanthine (11 μM), A2B receptor inhibitor MRS 1754 (1.97 nM), A3 receptor inhibitor MRS 1191 (92 nM) or A1 receptor agonist 2-Chloro-N6-Cyclopentyl adenosine (0.8 nM) for 30 minutes prior to adding pre-opsonized bacteria. The above concentrations correspond to the approximate Ki for each receptor targeting reagent. Reagents were purchased from Sigma. Percent killing was determined by dribble plating on blood agar plates and calculated in comparison to no PMN control under the exact same conditions (+/− treatments).

### Cell Death Assay

PMNs were incubated in HBSS/0.1% gelatin for 5 minutes at 37° C. The percentage of apoptotic and necrotic cells were then determined by flow cytometry using the FITC Annexin V apoptosis detection kit with PI (BioLegend) following manufacturer’s instructions.

### Bacterial Intracellular Survival Assay

To determine intracellular killing by PMNs, we performed a gentamicin-protection assay as previously described (Siwapornchai et al., 2020). Briefly, PMNs were infected at a multiplicity of infection (MOI) of 25 for 15 minutes at 37°C with bacteria pre-opsonized with homologous sera. Gentamicin (100ug/ml) was then added for 30 minutes to kill extracellular bacteria. The reactions were washed three times with HBSS and resuspended in 100ul HBSS/0.1% gelatin. To measure bacterial survival, the reactions were diluted and plated on blood agar plates and the percentage of the input inoculum that survived was calculated.

### Adenosine measurement

2×10^5^ bone marrow PMNs were incubated in OPH assay buffer for 5 minutes or infected with pre-opsonized *S. pneumoniae* TIGR4 for 45 minutes at 37° C. Adenosine level in the supernatants was measured using the Adenosine Assay Fluorometric Kit (Bio-Vision) as per manufacturer’s instructions.

### Statistics

All statistical analysis was performed using Prism8 (Graph Pad). CFU data were log-transformed to normalize distribution. Bar graphs represent the mean values +/− SD. Significant differences were determined by 1-sample t-test, Student’s t-test or one-way ANOVA followed by Dunnet’s or Tukey’s multiple comparisons test as indicated. All *p* values less than 0.05 were considered significant (as indicated by asterisks).

## Acknowledgements

We would like to thank Chelsie Armbruster for critical reading and discussion of the manuscript and John Leong for sharing reagents. This work was in part supported by National Institute of Health grant R00AG051784 and a University at Buffalo Clinical and Translational Science Institute pilot award CTSA1153519 to ENBG.

## Conflict of Interest

The authors declare no conflict of interest

## Author Contributions

MB conducted research, analyzed data and wrote paper. SS, AA and SEH conducted research and analyzed data. SER conducted research. ENBG designed research, wrote the paper and had responsibility for final content. All authors read and approved the final manuscript.

## Supporting Information Listing

### Supporting Experimental Procedures

#### ROS Assay

Intracellular and extracellular ROS production by PMNs was measured as previously described (Siwapornchai et al., 2020). Briefly, isolated PMNs were re-suspended in HBSS (Ca^2+^ and Mg^2+^ free) and acclimated at room temperature for one hour. The cells were then re-suspended in KRP buffer (Phosphate buffered saline with 5mM glucose, 1mM CaCl_2_ and 1mM MgSO_4_) and equilibrated at room temperature for another 30 minutes. The cells were seeded in 96-well white LUMITRAC™ plates (Greiner Bio-One) at 5×10^5^ PMNs per well, treated either with pre-opsonized *S. pneumoniae* TIGR4 at a MOI of 25, 100 nM Phorbol 12-myristate 13-acetate (PMA) (Sigma) were used as a positive control or mock treated with buffer containing 3% mouse sera (uninfected). For detection of extracellular ROS, 50μM Isoluminol (Sigma) plus 10U/ml HRP (Sigma) were added, while for detection of intracellular ROS, 50μM Luminol (Sigma) was added to the corresponding wells. Luminescence was immediately read over a period of one hour at 37° C in a pre-warmed Biotek Plate reader. Wells with buffer and Isoluminol plus HRP or Luminol alone were used as blanks.

#### Generation of GFP-expressing *S. pneumoniae*

AC316-GFP was based on a GFP construct in *S. pneumoniae* D39, where a GFP-antibiotic resistance cassette transcriptional fusion was translationally fused to the end of *hlpA*, with a flexible linker connecting the two pieces (Kjos et al., 2015). The transcriptional fusion of sfgfp-SpectinomycinR was translationally fused to the end of the *hlpA* gene by removing the stop codon and placing a flexible linker with a GSGGEAAAKG motif (Arai, Ueda, Kitayama, Kamiya, & Nagamune, 2001). The individual pieces of DNA were designed with overlapping sequences such that they could be assembled into one contiguous piece of DNA. The *hlpA* gene and upstream AC316 sequences were amplified using primers hlpA-up-F and hlpA-link-R (Table 1). The sfgfp gene was amplified from plasmid pTHSSd_34 using primers gfp-link-F and gfp-R-spec (Table 1). pTHSSd_34 was a gift from Christopher Voigt (Addgene plasmid # 59960). The *aad* gene encoding spectinomycin resistance was amplified from pMagellan6 using the primers spec-F-gfp and spec-R-hlpA (Table 1). The region downstream of *hlpA* was amplified with primers hlpA-down-F-spec and hlpA-down-R (Table 1). The four linear pieces of DNA were mixed in equimolar amounts, and splicing by overlap extension PCR was used to assemble one linear piece of DNA, which was amplified using the primers hlpA-up-F and hlpA-down-R (Table 1). The resulting linear DNA was transformed into AC316 as previously described (Siwapornchai et al., 2020), and transformants were screened for fluorescence.

#### Antimicrobials ELISA

1×10^6^ Bone marrow PMNs were infected with pre-opsonized *S. pneumoniae* TIGR4 at a MOI of 2 for 45 minutes at 37° C. The cell free supernatants were then collected and assayed for Mouse MPO levels (Invitrogen) and CRAMP (MyBiosource) levels by ELISA as per manufacturer’s instructions.

#### Bacterial Uptake Assay

PMNs were infected with GFP-expressing *S. pneumoniae* at a MOI of 10 or mock-treated in 100μl reactions of HBSS/0.1% gelatin. Reactions were rotated for 15 minutes at 37°C. To differentiate between associated vs. engulfed bacteria, cells were then washed and resuspend in FACS buffer (HBSS/1% FBS). The cells were and stained with primary rabbit polyclonal anti *S. pneumoniae* capsular serotype 4 antibodies (Cederlane) followed by secondary PE-conjugated anti-Rabbit IgG (12473981; Invitrogen). Cells were analyzed by flow cytometry to determine the percentage of PMNs that associated with bacteria (GFP^+^ PMNs). GFP^+^ cells were gated on and the percentage of engulfed bacteria (GFP^+^/ PE^−^) vs. extracellular bacteria (GFP^+^PE^+^) was determined.

#### Western Blots

PMNs were solubilized in radioimmunoprecipitation assay buffer (1% Triton X-100, 0.25% sodium deoxycholate, 0.05% SDS, 50 mM Tris-HCl [pH 7.5], 2 mM EDTA, 150 mM NaCl, 1 mM phenylmethylsulfonyl fluoride, 1 mM sodium orthovanadate, 10 mg/liter each of aprotinin and leupeptin). Protein concentrations of the lysate supernatants were quantified using bicinchoninic acid kit (Pierce). Equal quantities of each sample were run on Mini-PROTEAN TGX Stain-Free Precast Gels (BioRad) and transferred to polyvinylidene difluoride (PVDF) membranes. Incubations with primary antibodies at 1:1000 dilutions were done overnight at 4°C. Incubation with secondary antibodies (1: 5000 dilutions) coupled to horseradish peroxidase was done for 1 hour at room temperature. Clarity Western ECL Substrate (BioRad) was used for detection of all blots using the ChemiDoc XRS+ system (BioRad). For loading controls, blots were treated stripping buffer containing glycine, Triton-X 100 and SDS, and probed for GAPDH. Primary antibodies against A1 (ab82477) were purchased from Abcam. Primary antibodies against GAPDH (MA5-15738) and horseradish peroxidase-conjugated secondary antibodies (31460 and 31430) were purchased from Invitrogen. Images were quantified using ImageJ (version 1.51e) software. Band densities were measured, and the background was subtracted. Background-corrected densities of A1 receptor proteins were normalized to GAPDH.

### Supporting Figure Legends

**Supporting Figure 1.**
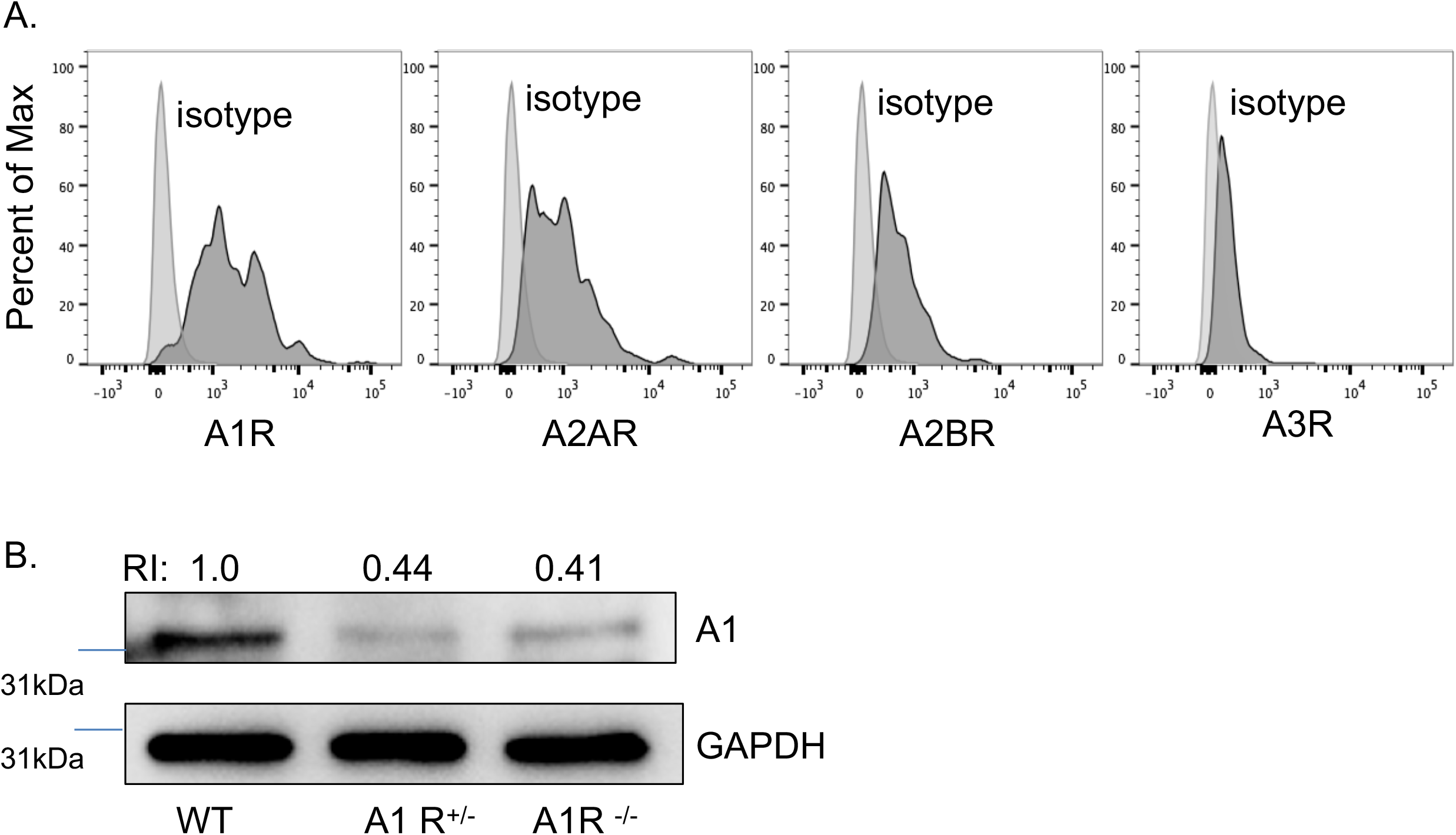
Expression of adenosine receptors on PMNs. Bone marrow PMNs were isolated from young C57BL/6 mice and expression of adenosine receptors was assessed by flow cytometry. Histograms shown are representative data from one of three separate experiments.

**Supporting Figure 2.**
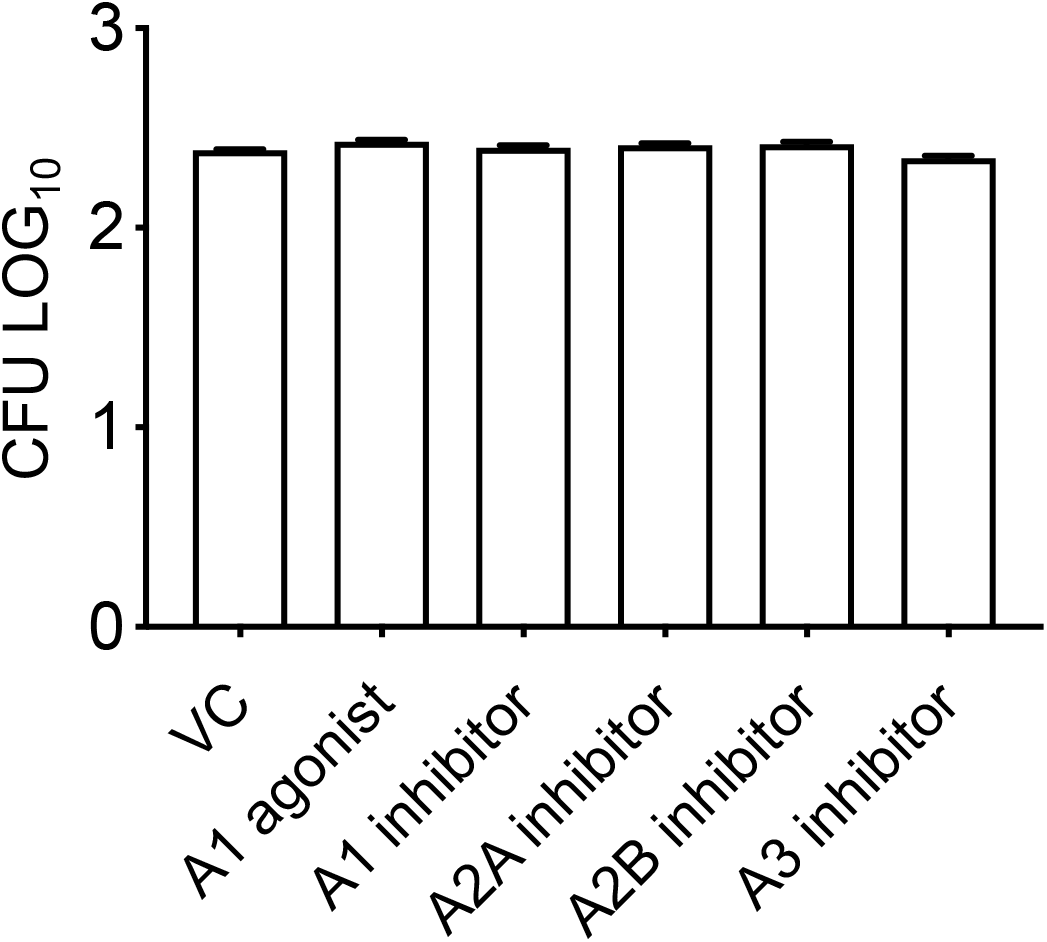
Effect of adenosine pathway drugs on bacterial viability. *S. pneumoniae* TIGR4 were treated with PBS control (VC), A1 receptor agonist 2-Chloro-N6-cyclopentyladenosine (0.8 nM), A1 receptor inhibitor 8-Cyclopentyl-1-3-dipropylxanthine (3.9nM), A2A receptor inhibitor 3,7-Dimethyl 1-1-propargylxanthine (11 μM), A2B receptor inhibitor MRS 1754 (1.97nM) or A3 receptor inhibitor MRS 1191 (92 nM) for 40 minutes. The number of viable bacteria was determined by plating on blood agar plates. Data shown are representative from one of three separate experiments where each condition was tested in triplicate.

**Supporting Figure 3.**
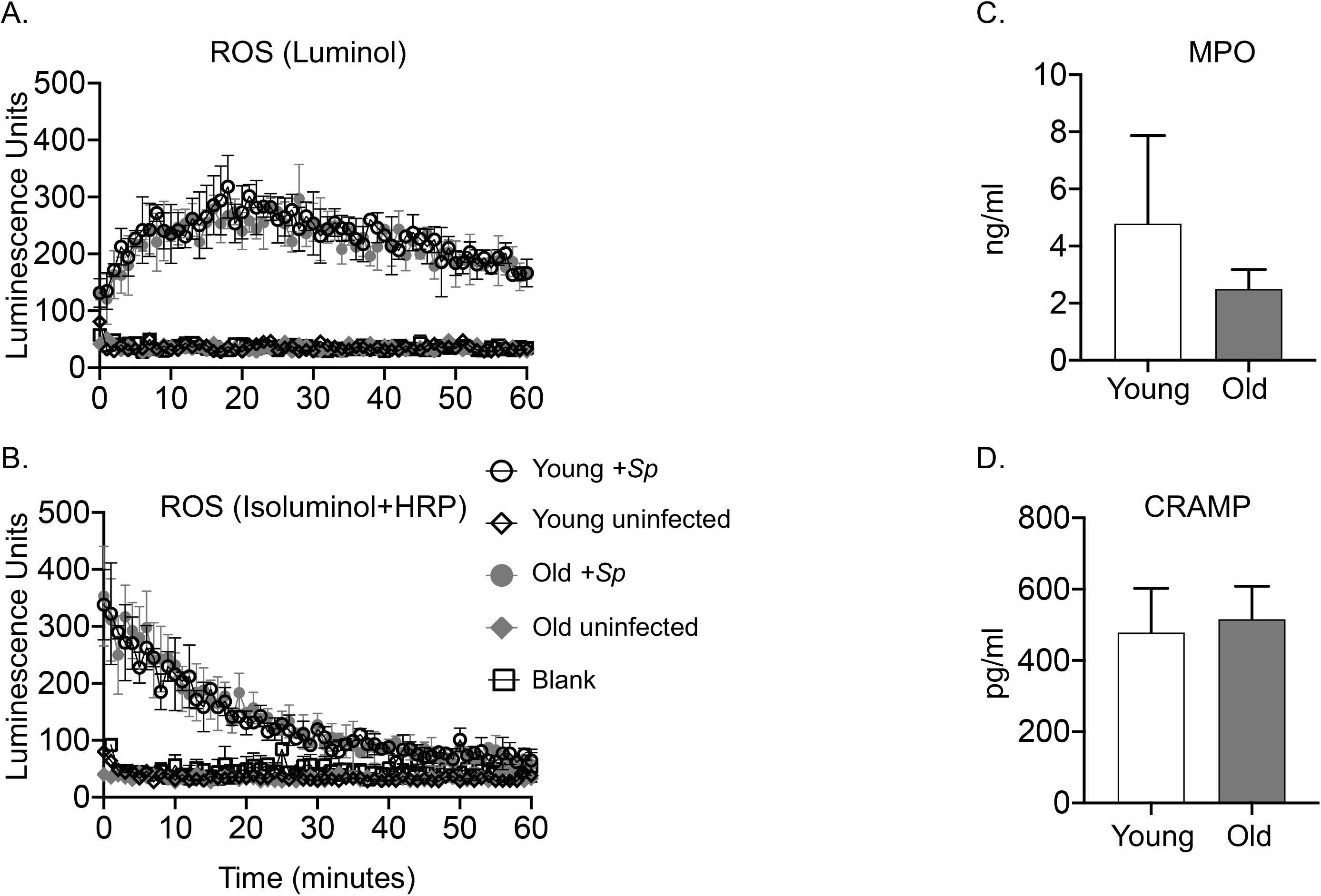
Aging does not affect ROS production by PMNs in response to *S. pneumoniae.* PMNs were isolated from the bone marrow of young (2 month) and old (18-22 month) male C57BL/6 mice. PMNs were then infected with *S. pneumoniae* pre-opsonized with homologous sera (+*Sp*) or treated with 3% matching sera (uninfected). Intracellular ROS production measured by chemiluminescence of Luminol (A) and extracellular ROS production measured by chemiluminescence of Isoluminol in the presence of HRP (B). (A-B) Representative data are shown from one of three separate experiments with one mouse per strain per experiment where each condition is tested in triplicates. (C-D) PMNs from the indicated mice were incubated for 45 minutes at 37°C with pre-opsonized *S. pneumoniae.* The supernatants were collected and assayed for CRAMP and MPO levels. Data were pooled from three experiments.

**Supporting Figure 4.**
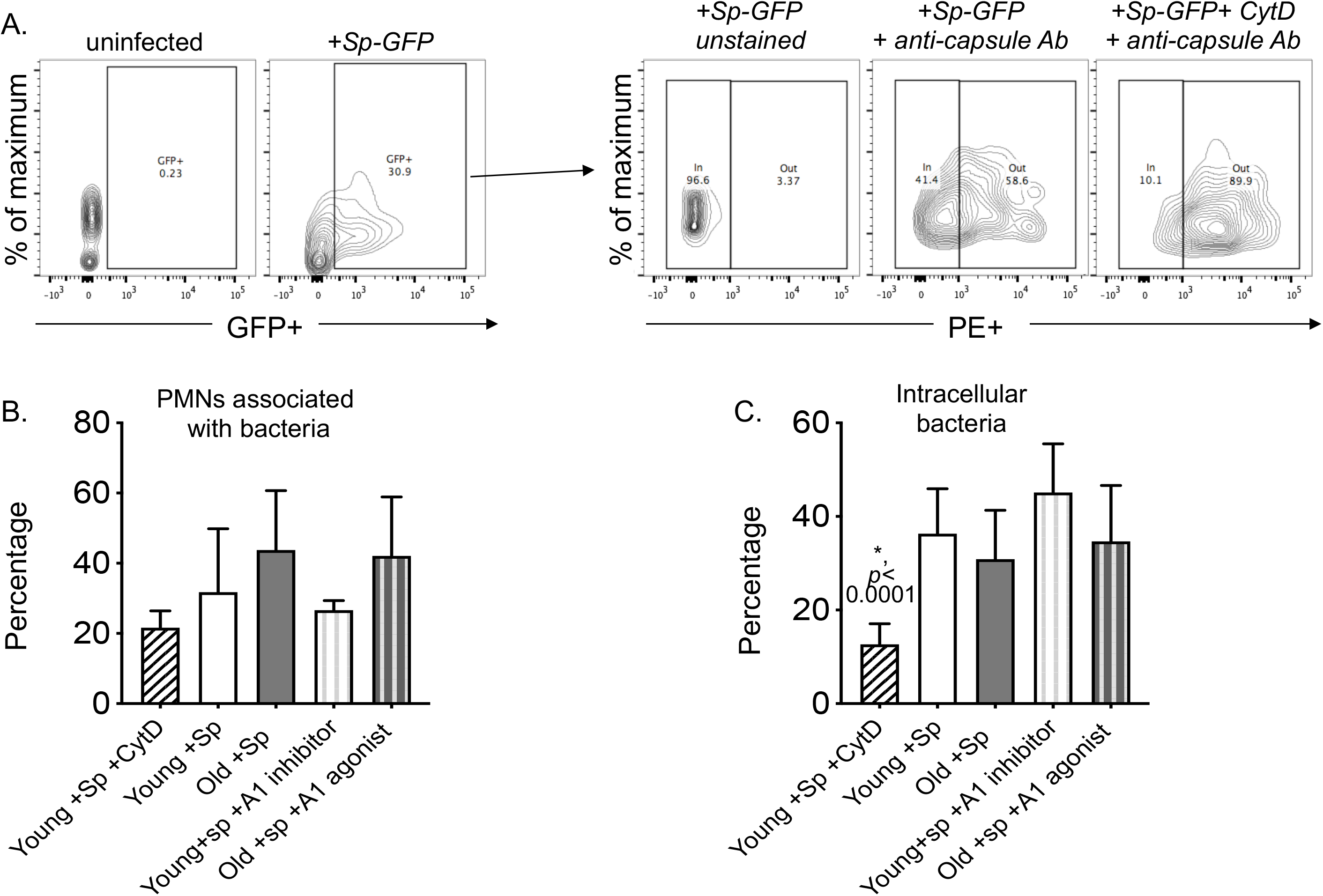
Aging does not affect bacterial uptake by PMNs. PMNs isolated from the bone marrow of young and old C57BL/6 mice were left untreated or treated with Cytochalasin D (CytD 20μM), A1 receptor agonist 2-Chloro-N6-cyclopentyladenosine (0.8 nM) or A1 receptor inhibitor 8-Cyclopentyl-1-3-dipropylxanthine (3.9nM) where indicated. PMNs were then infected with GFP-expressing *S. pneumoniae* at a MOI of 10 or mock-treated for 15 minutes and then stained with primary unconjugated anti-*S. pneumoniae* capsular serotype 4 antibodies followed by PE-conjugated secondary antibodies. Cells were analyzed flow cytometry to determine the percentage of PMNs that associated with bacteria (GFP^+^ PMNs). GFP^+^ cells were gated on and the percentage of engulfed bacteria (GFP^+^/ PE^−^) vs. extracellular bacteria (GFP^+^PE^+^) was determined. (A) Density plots shown are representative data from young mice and bar graphs are quantification of (B) the percentage of PMNs associated with bacteria and (C) the percentage of associated bacteria that was engulfed by PMNs. Data shown are pooled from three separate experiments (n=3 biological replicates or mice per strain) where each condition was tested in triplicate (n=3 technical replicates) per experiment.

## Notes

### Competing Interest Statement

The authors have declared no competing interest.

